# Adulthood Deficiency of the Insulin-like Growth Factor-1 Receptor in Hippocampal Neurons Impairs Cell Structure and Spatial Learning and Memory in Male and Not Female Mice

**DOI:** 10.1101/2021.08.08.455596

**Authors:** Cellas A. Hayes, Erik L. Hodges, Jessica P. Marshall, Sreemathi Logan, Julie A. Farley, Daniel B. Owens, William E. Sonntag, Nicole M. Ashpole

**Affiliations:** Department of BioMolecular Sciences, University of Mississippi School of Pharmacy, University of Mississippi, University, MS 38667; Department of Rehabilitation Sciences, University of Oklahoma Health Sciences Center, OK; Center for Geroscience and Healthy Brain Aging, Department of Biochemistry and Molecular Biology, University of Oklahoma Health Sciences Center, OK; Research Institute of Pharmaceutical Sciences, University of Mississippi School of Pharmacy, University of Mississippi, University, MS 38677

**Keywords:** IGF-1, somatomedin C, cognition, sex-specific, neuronal, hippocampus

## Abstract

Reductions in insulin-like growth factor-1 (IGF-1) are associated with cognitive impairment and increased risk of neurodegenerative disease in advanced age. In mouse models, reduced IGF-1 early-in-life leads to memory impairments and synaptic dysfunction; however, these models are limited by systemic reductions in IGF-1. We hypothesized that IGF-1 continues to promote hippocampal neuron structure and function after development, and as such, the loss of IGF-1 signaling in adult neurons would lead to impaired spatial learning and memory. To test this, the IGF-1 receptor (IGF-1R) was genetically targeted in hippocampal neurons of adult male and female mice. Male mice deficient in neuronal IGF-1R exhibited spatial learning impairments as evidenced by increased pathlength and errors in the radial arm water maze. No differences in learning and memory were observed in female mice. Golgi-Cox staining revealed a reduced number of dendritic boutons of neurons the CA1 region of the hippocampus in male mice. Decreased MAPK and increased ROCK activity were also observed in these tissues. In vitro studies revealed that impaired neurite outgrowth due to inhibited IGF-1R signaling could be rescued by pharmacological inhibitors of ROCK. However, ROCK inhibition in neuronal IGF-1R-deficient mice did not fully rescue learning impairments or bouton numbers. Together, our study highlights that IGF-1 continues to support spatial learning and memory and neuronal structure in adulthood.

## INTRODUCTION

Insulin-like growth factor-1 (IGF-1) promotes of brain development and pubertal growth. While much focus is placed on the impact of IGF-1 during the developmental phases of life, the growth hormone (GH) and IGF-1 signaling axis are implicated in critical behavioral and cellular functions throughout the lifespan. In fact, one of the hallmarks of aging is the reduction in the GH/IGF-1 signaling (Ashpole et al., 2015b; Lopez-Otin et al., 2013). Circulating IGF-1 levels are reduced 30-60% in elderly human and animal models (Florini et al., 1985; Johanson and Blizzard, 1981; Rudman et al., 1981). This decrease is temporally associated with the onset of cognitive decline and increased risk of neurodegenerative diseases and dementias (Arwert et al., 2006; Deijen et al., 2011; Frater et al., 2018; Sonntag et al., 2005b; Trejo et al., 2007). Importantly, supplementation of IGF-1 has been shown to reverse cognitive impairment in aged animals (Markowska et al., 1998), suggesting that maintenance of the IGF-1 signaling cascade remains important for neuronal function after development. Recombinant IGF-1 is FDA approved for the treatment of early-life growth deficiencies, however its use is limited to this developmental growth window and has not been tested in cognitively-impaired, elderly humans. As an anabolic hormone associated with increased cancer risk, systemic administration of exogenous IGF-1 is not an ideal therapeutic for adult and elderly individuals (Shanmugalingam et al., 2016; Sonntag et al., 2005a). Instead, it may be advantageous to selectively target one of the downstream signal transduction cascades that are dysregulated when IGF-1 levels decline in advanced age. In order to do this, a better mechanistic understanding of the influence of IGF-1 on cognitive behaviors in adulthood is needed.

Interruptions in the IGF-1 signaling cascades lead to significant impairments in long-term potentiation and learning and memory (Cao et al., 2011; D’Mello et al., 1997; Gazit et al., 2016; Glasper et al., 2010; Maher et al., 2006). In the developing brain, the beneficial effects of IGF-1 are thought to be mediated by pro-survival and pro-growth pathways. Activation of the IGF-1 receptor (IGF-1R) and the downstream PI3K, Akt, and MAPK signaling cascades enhance neuronal survival, growth, and excitability (Cheng et al., 2003; Scolnick et al., 2008; Trejo et al., 2007). Little is known about the effects of IGF-1 on growth-limiting pathways. In fact, several intracellular signaling cascades limit neuronal structure throughout the lifespan, including the RhoA/Rho-associated coiled-coil containing protein kinase (ROCK) pathway, which stabilizes neurofilament rearrangement (Bito et al., 2000; Hirose et al., 1998). Increased RhoA/ROCK is associated with spine retraction, synapse elimination, decreased neuronal excitability, and impaired learning and memory (Dash et al., 2004; Lamprecht et al., 2002; Sunico et al., 2010). Expression and activity of the RhoA/ROCK pathway are upregulated in advanced age and are further upregulated in cognitively-impaired aged animals (VanGuilder Starkey et al., 2013). In aged rats and mouse models of Alzheimer’s Disease and intellectual disability, ROCK inhibition rescues spatial learning and memory impairments (Busti et al., 2020; Guo et al., 2020; Huentelman et al., 2009; Meziane et al., 2016; Yu et al., 2017). If IGF-1 regulates ROCK activity, then the age-related loss of IGF-1 could account for the upregulation of this growth-limiting pathway.

Given the temporal associations with reduced circulating IGF-1, increased ROCK, and impaired cognition, we hypothesized that the loss of IGF-1 in adulthood leads to decreased growth-promoting signals as well as increased growth-restricting signals, which ultimately causes loss of neuronal structure inherently leading to learning impairments. The common mammalian model organisms in which IGF-1 signaling is altered involve reducing hepatic production of IGF-1, which is responsible for releasing IGF-1 into circulation. This alters systemic IGF-1 signaling, thereby confounding our understanding of the cell-specific activities. Indeed, reductions in circulating IGF-1 alters the structure and function of several cell types, including astrocytes, microglia, and the cerebrovasculature (Grinberg et al., 2013; Logan et al., 2018; Park et al., 2011; Prabhu et al., 2019; Tarantini et al., 2016a). Most IGF-1-deficient animal models also have reduced IGF-1 levels throughout both development and adulthood, which may instigate compensatory effects that confound interpretation. To date, studies have not delineated whether IGF-1 continues to regulate learning and memory by acting specifically on neurons after development. To address this, we utilized genetic tools to specifically target IGF-1R signaling in hippocampal neurons in adulthood and then assessed the molecular, cellular, and behavioral changes induced in the months following knock-out.

## MATERALS AND METHODS

### Animals

All procedures were approved by the Institutional Animal Use and Care Committees of the University of Mississippi and the University of Oklahoma Health Sciences Center. IGFR ^*f/f*^ were purchased from Jackson laboratories (B6;129-Igf1rtm2Arge/J) and bred in house to develop experimental cohorts (N=128 experimental mice). Spatial learning and memory was assessed at both research sites to verify reproducibility following lab relocation. These mice express Lox P sites surrounding exon 3, which encodes a key region of the ligand-binding domain of IGF-1R. All mice were genotyped to verify *igfr*^*f/f*^, using the primers and PCR protocols designed by Jackson laboratories, as previously reported (Prabhu et al., 2019). At the University of Oklahoma Health Sciences Center, mice were housed under specific pathogen-free barrier conditions in the Rodent Barrier Facility under a controlled photoperiod (12 h light; 12hdark) with unlimited access to water and standard AIN-93G diet food (ad libitum). At the University of Mississippi, mice were housed in ventilated racks (Lab Products, Inc. RAIR Envirogaurd ventilated cages) with enriched bedding under a 12:12 hour photoperiod, with 5053 Pico Lab chow and water ad libitum. Block randomization was utilized to ensure balanced group size and treatment groups were separated by cages. Male and female cohorts were treated independently. Following behavioral analysis, mice were euthanized via CO^2^ asphyxiation for ex vivo tissue isolation and analysis.

Maintenance of *igfr*^*f/f*^ homozygosity was verified in all offspring after weaning (3-5 mice per cage). Following 2-3mm tail removal, DNA was extracted using the RED Extract-N-Amp Tissue PCR XNAT, following manufacturer’s instructions (Sigma). The recommended IGFR flox primers and PCR protocols designed by Jackson laboratories were utilized (primers purchased from Sigma). PCR products were run on a 1.5% agarose gel stained with green glow (Denville) at 120V in 1XTB for 1 hour. UV imaging identified PCR products. All gels contained a 100bp DNA ladder (ThermoFisher), an IGFR ^*f/f*^ control lane, and an IGFR^+/+^ control lane.

Primary neuron cultures were developed using tissue from post-natal (P1) *igfr*^*f/f*^ mice. Male and female pups were not separated during isolation. Neuron cultures were also developed using tissue from embryonic (E-19) and post-natal (P1) Sprague Dawley rats. Timed pregnant rat dams were purchased from Envigo and were euthanized along with the pups.

### Hippocampal Adeno-Associated Virus Injections

At three months of age, cohorts of male and female *igfr*^*fl/fl*^ mice received stereotaxic injections of AAV9-Syn-Cre or control AAV9-Syn-GFP (purchased purified virus from Penn Viral Vector Core) in the dentate gyrus and CA1 subregions of the hippocampus (Barbash et al., 2013; Stoica et al., 2013). Mice were anesthetized with ketamine/xylazine or with 5% isoflurane and the anesthetic plane was verified with a toe pinch. Mice were placed on a heating pad, EMLA cream (Lidocaine (2.5%) and Prilocaine Cream (2.5%), USP) was applied to the stereotactic ear bars, and a dab of Puralube was placed on each eye. Saline (1mL) was subcutaneously injected to prevent dehydration. The topical area of the cranium was scrubbed with 1% chlorohexidine or 70% ethanol and the hair was trimmed. A 5-8mm incision exposed the sagittal suture, the skull was leveled, and bregma and lambda measurements were used to determine injection coordinates (posterior -0.28mm, medial/lateral +/- 0.25). A high speed microdrill was used to puncture the skull at the calculated position and a Hamilton syringe was guided to the injection depth (−0.27 for CA1 and - 0.3mm for DG). Bilateral injections were performed in all mice with each hippocampal hemisphere receiving a total of 1μl diluted virus (0.5μl per injection depth, administered at 0.1μls/10 seconds). Bone wax (Surgical Specialties) was applied to the injection holes and the incision was then mattress sutured (Nylon sutures). Meloxicam or Bupivicaine (Henry Shrien) was administered for pain (immediately and as needed with 48 hours of procedure).

### Hydroxyfasudil Osmotic Pump Implantation

A cohort of mice received hydroxyfasudil (Sigma) or control DMSO/ddH2O for 14 days via implanted subcutaneous mini osmotic pumps (Alzet). Pumps were equilibrated in 0.9% saline overnight and loaded using the manufacturer’s instructions. Mice were weighed to determine 10mg/kg dosing of hydroxyfasudil (diluted in 18% DMSO and 82% ddH2O) in 200μl buffer across 14 days. Groups were block randomized to receive hydroxyfasudil or vehicle control; however, 6 controls (*IGFR-WT) did not receive pumps (all remaining mice within that experimental cohort did). Mice were anesthetized using 5% isoflurane and the osmotic pump was inserted under the skin at the nape of the neck. The incision site was mattress sutured and lidocaine was administered topically on the wound. Pumps remained in place throughout the behavioral assessment. No adverse effects were noted.

### Quantitative PCR

Total RNA was isolated using the RNAeasy mini kit (Qiagen) and converted to cDNA using High-Capacity RNA-to-cDNA kit (Applied Biosystems). RT-PCR was performed using TaqMan Universal PCR Master Mis (Life Technologies) and TaqMan validated primers on the CFX Connect Real-time PCR detection system (Bio-Rad). Results were normalized to housekeeping genes, HPRT and B2M. The experimenter was blinded to treatment group during experimentation, and the assignment code was revealed for delta-delta CT analysis.

### Radial Arm Water Maze

Two months following viral injection, spatial learning and memory was assessed in a flooded -eight-arm maze with a submerged escape platform and spatial clues. Water (5-7cm depth) was mixed with white food coloring to create an opaque color to hide the safety platform. Mice were acclimated to the escape platform before beginning the acquisition phase of the maze, which consisted of 24 training trials (8 trials per day for 3 consecutive days). On day 10, memory was assessed in a 60-second probe trial. The platform was then moved to an alternate arm for a reversal phase where extinction and re-learning were observed for an additional 8 trials. Total pathlength to escape platform, latency to escape, the number of errors made, and velocity were calculated using Ethovision Software (Noldus Information Technology Inc., Leesbur, VA, USA). Six male mice that did not undergo stereotaxic AAV injection and were included as an additional treatment control group in this maze.

### Novel Object Relocation and Recognition

Assessment of working memory with the novel object (NO) and novel location (NL) assay was carried out over 5 exploration trials in a white-walled open arena containing 2 objects. Each trial was 5 minutes in length, with a 5-minute intertrial interval. Trials 1, 2, and 3 were used to familiarize the mouse with the arena. In trial 4, the object was relocated (NL) and in trial 5, the object was replaced (NO). The time spent at each object was used to calculate the discrimination index for NL (time spent at NL/(time spent at NL + time spent at F3)) and NO (time spent at NO/(time spent at NO + time spent at F3)) using Ethovision Software (Noldus Information Technology Inc.).

### Western Blot

Hippocampal tissue and primary neuron cultures were lysed in RIPA buffer (Fisher) with protease and phosphatase inhibitors (Calbiochem, Roche). Protein concentration in the lysates was calculated using an RC/DC Assay, per manufacturer’s recommendations (BioRad). Protein (10-20μg) was denatured and loaded into 4-12% precast bis-tris gels (Invitrogen, Thermo Fisher). Following electrophoresis, gels were transferred to nitrocellulose membranes, blocked in 5% BSA in 1XTBST, incubated with primary antibody, washed, and incubated with secondary antibodies. All antibodies, dilutions, and RRIDs are described in **Supplemental Table 1**. Blots were imaged and analyzed on the Licor Odyssey CLx.

### Golgi Staining

A sagittal hemisphere of the brain was stained using the FD Rapid Golgi Stain Kit (FD Neuro Technologies, Inc.), following the manufacturer’s recommendations. Due to equipment procurement between studies, female specimens were sliced at 100μm on a cryostat (Leica), while male specimens were sliced at 250μm after being solidified in 3% agarose in ddH20 on a vibratome (Precision Instruments) with an oscillation speed of 6.9 and speed of 1.9. Brightfield imaging was performed on a Nikon Ti2-E inverted microscope with a 60x objective. Z-stacked images were acquired and a 20μm line was drawn on dendrites selected at random to quantify the number of boutons across a controlled length. Two dendrites per field were selected by a blinded observer. Images were taken of both sub-regions, DG and CA1, for separate quantification.

### Primary neuron cultures

Hippocampal and/or mixed hippocampal/cortical tissue was isolated from *igfr*^*f/f*^ mice or wild-type Sprague Dawley rats to establish primary cultures of neurons, following our previously established protocols (Ashpole and Hudmon, 2011). Tissue was enzymatically digested with Papain (Worthington Biochemical Corporation) at 37°C for 20-30 minutes, mechanically triturated, and filtered. Cells were pelleted and resuspended in fresh neuronal plating media Neurobasal media (Life Technologies), 5% FBS (Cellgro or Sigma), 1X Glutamax (Life Technologies), and 1X N2 Supplement (Life Technologies). The next day, media is changed to neuronal growth media (Neurobasal media, 1X B27 (Life Technologies), 1X Penicillin/ Streptomycin, and 1X Glutamax), and is maintained in this media with mitotic inhibitors until experimental use. For experiments were neurons where IGF-1R is genetically reduced, AAV9-Syn-Cre and control AAV9-Syn-GFP was transduced at day 5 of culturing. For all pharmacological treatments, drugs were diluted in fresh media and administered on days 8-10 of culture, and experimental endpoints were 24 hours later.

### Sholl analysis

Hippocampal neurons were grown on coverslips and transfected with CMV-tdTomato (Addgene) between days 2 and 5 of culture. Treatments were administered between days 8-10 and 24 hours later, neurons were fixed in 4% paraformaldehyde in phosphate buffer, pH 8.0. Coverslips were washed with 1XPBS and mounted using Prolong Gold Antifade with DAPI. Transfected neurons were selected at random by a blinded observer and imaged using a Nikon Ti2-E inverted microscope with a 20x objective and 540/70 excitation/emission filter. Multiple fields (6-10) were selected at random on each coverslip (8-10) for analysis with Neurolucida (MBF Biosciences) or ImageJ (Fuji) Sholl plug-ins. The average area under the curve for each coverslip was interpolated using SigmaPlot v13 (Systat).

### ROCK Activity Assay

Hippocampal tissue or 10cm dishes of primary mixed cortical/hippocampal neurons were lysed in RIPA buffer (Fisher) containing 1X protease and phosphatase inhibitors (Calbiochem, Perkin Elmer) for use in the Rho-associated Kinase Activity Assay (Millipore). The manufacturer’s recommendations were followed for all steps of the assay. This assay does not differentiate ROCK-1 and ROCK-2 activity.

### RhoA Activation Assay

GTP-bound RhoA was quantified using the RhoA Pull-down Activation Assay Kit (Cytoskeleton, Inc), following the manufacturer’s recommendations. Mixed neuronal cultures were treated with pharmacological inhibitors for 24 hours and subsequently lysed in the proprietary lysis buffer. GST-tagged Rhotekin-RBD beads were incubated in the lysate prior to washing, eluting, and western blotting with the materials provided in the kit.

### Experimental Design and Statistical Analysis

All analysis and graph generation was performed using Sigma Plot v13 software and/or R Version 64 3.6.1 in addition, to RStudio with package lmerTest version 3.1-0. For comparisons of two treatment groups (WT vs KO or Vehicle v PPP), a Students unpaired t-test was utilized and normality was assumed. To compare concentration-dependence, a one-way ANOVA with posthoc Dunnett’s test vs vehicle control was performed. Many experiments were set as 2×2 factor designs (genetic treatment and pharmacological treatment), thus two-way ANOVA with posthoc Fishers LSD or Bonferonni were utilized. Behavioral assays with multiple trials were set as two-way repeated-measures ANOVAs. Male and female cohorts were independently assessed and direct comparisons were not performed. Behavioral experiments were constructed to detect a 20% difference between treatment groups, with a power of 0.8 using a type 1 error rate of 0.05. Based on preliminary data, we estimated n=8 mice were needed per treatment group. All data were expressed with mean +/- standard error and p<0.05 was used to define statistical significance. The statistical comparisons that accompany our experimental design and working hypotheses used are defined in Figure Legends. All p values and final sample sizes can be found in **Supplemental Table 2**.

All mice and neuron images were coded to blind experimenter to treatment groups during testing. Neurons were selected at random for structural studies, with multiple cells per brain slice or coverslip assessed for the rigorous representation of the data. No mice were excluded from the behavioral and/or molecular studies; however, 3 mice did die during hippocampal injection surgery.

## RESULTS

### Learning impairments with neuronal IGFR deficiency

Hippocampal neuron-specific IGF-1R deletion (*IGFR-KO) was induced in male and female *igfr*^*f/f*^ mice at 3-4 months of age via stereotaxic injections of Cre recombinase encoding viral vectors (AAV9-Syn-Cre or control AAV9-Syn-GFP) (**Figure 1A**). mRNA expression of IGFR, IGF-1, and other growth-promoting signals was quantified in the dorsal hippocampus to verify specific reductions in the floxed region of IGF-1R exon 3. IGF-1R was reduced by 39.6% in these neuron-specific *IGFR-KO male and female mice, compared to GFP-transduced controls (*IGFR-WT) (**Figure 1B**, p=0.033, n=9). No significant differences in the expression of growth hormone receptor (GHR), insulin receptor (InsR), or IGF-1 were observed in the hippocampi of *IGFR-KO mice (**Figure 1B**, p>0.05, n=9). Details of exact p-value comparisons throughout the entire result section are described in **Supplemental Table 1**.

**Figure 1:**
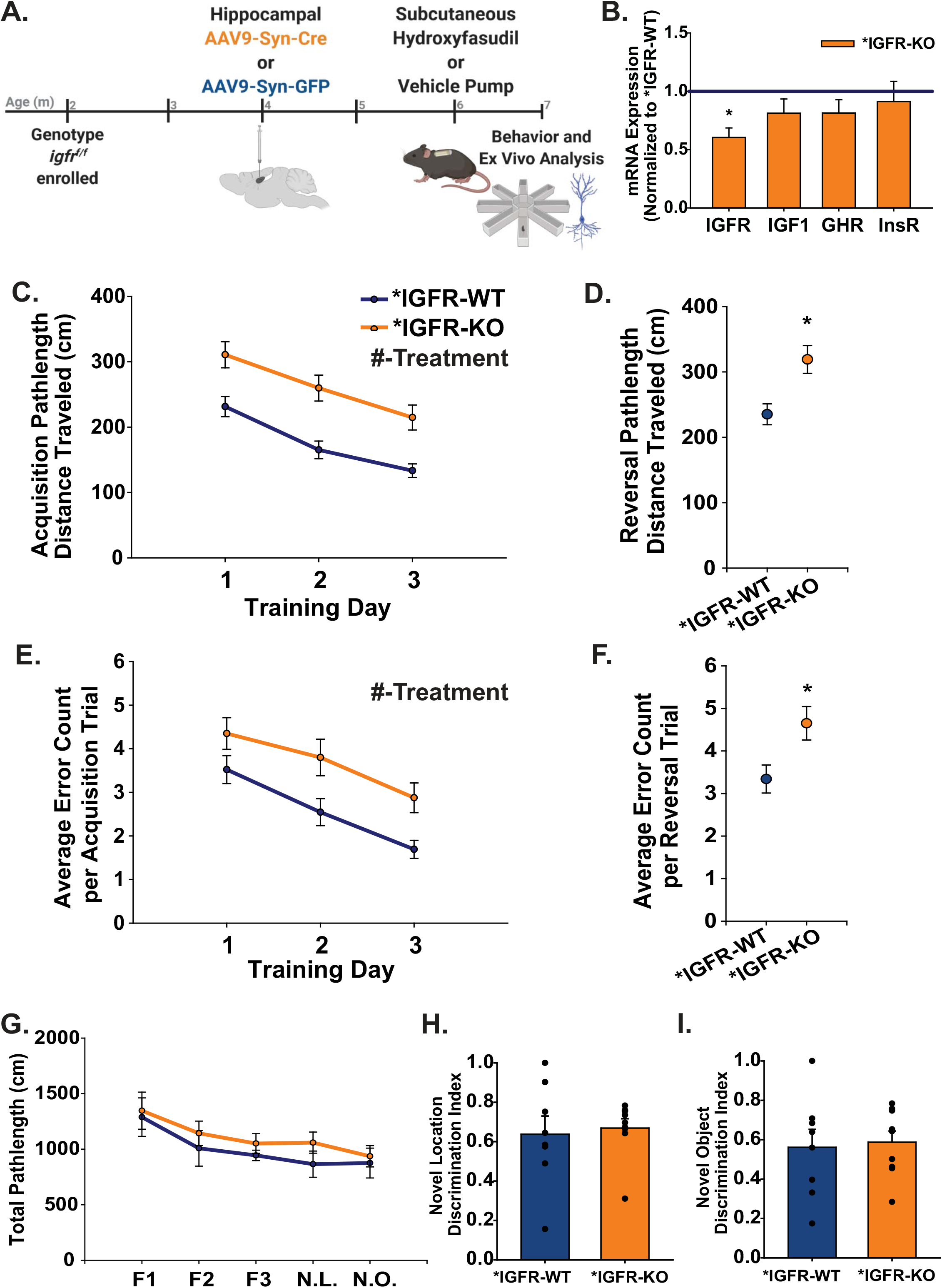
Adult IGF-1R Signaling in Neurons Regulates Spatial Learning and Memory in Male Mice. **(A)** Experimental design and timeline of hippocampal neuron-specific deletion of IGFR (*IGFR-KO), behavioral testing, and ex vivo analysis. Image made with biorender.com **(B)** Hippocampal mRNA expression of IGF-1R, IGF-1, growth hormone receptor (GHR), and insulin receptor (InsR) in *IGFR-KO mice, normalized to control *IGFR-WT (mean +/- SEM, n=9). Geometric means of HPRT and B2M were utilized as housekeeper genes. The asterisk indicates a significant difference of each gene vs*IGFR-WT control (*p<0.05, t-test). (**C-F**) Assessment of spatial learning and memory in male *IGFR-WT and *IGFR-KO mice within the radial arm water maze. Average pathlength traveled per trial by male mice during the acquisition phase **(C)** and reversal phase **(D)** of the radial arm water maze. Average number of errors per trial in the acquisition phase **(E)** and reversal phase **(F)** of the radial arm water maze. The # denotes p<0.05 across treatment groups in the acquisition phase (mean +/- SEM, n=10,11, two-wayANO VA repeated measures). The asterisk denotes p<0.05 vs *IGFR-WT in the reversal phase (t-test). **(G)** Average pathlength of male *IGFR-WT and *IGFR-KO mice in each of the five phases of the novel object/relocation task (p>0.05, two-way ANOVA repeated measures). (**H**) Average discrimination between the object relocated during the novel location trial (p>0.05, t-test). (**I**) Average discrimination between the new and old object during the novel object (p>0.05, t-test). All novel object data presented as mean +/- SEM, n=11,10.

Two to three months following knockout, spatial learning/memory was assessed in the radial arm water maze. *IGFR-WT and *IGFR-KO mice both learned to locate the escape platform over the three-day training phase, as indicated by a reduction in the total pathlength traveled to reach the escape platform (**Figure 1C**) and the total number of errors made in finding the platform (**Figure 1E**). However, *IGFR-KOs traveled significantly more and made more errors throughout the learning phase than controls (**Figure 1C**, p=0.0002; **Figure 1E**, p=0.018; n=10,11). When memory extinction and relearning was assessed in a reversal phase of the maze, male *IGFR-KO mice again traveled significantly more and made more errors than controls (**Figure 1D**, p=0.0017; **Figure 1F**, p=0.011, n=10,11). *IGFR-WT controls performed similarly to mice that had not undergone AAV injection, suggesting the surgical procedure alone did not alter spatial learning and memory (**Supplemental Figure 1A**)

Unlike males, female *IGFR-KO mice did not show significant impairments in spatial learning and memory compared to *IGFR-WT controls. Both groups of females exhibited similar reductions in pathlength and the number of errors over training days; no between-group differences were observed and no differences were noted in the reversal phase one week later (**Supplemental Figure 1B**, p>0.05, n=9,10).

In addition to testing spatial learning and memory, working memory was assessed in the novel object replacement and relocation task. No significant difference in exploration or novelty discrimination was observed in male *IGFR-KOs (**Figure 1G-I**, p>0.05, n=8,9) or female *IGFR-KOs; however, *IGF-1R KO females explored the arena significantly more than female *IGFR-WT females (**Supplemental Figure 1C**, p<0.01, n=10).

### Structure changes in Neurons Deficient in IGF-1R

Following behavioral analysis, the neuronal structure within the CA1 region of the hippocampus was visualized using Golgi-Cox staining (**Figure 2A-B**) and synaptic bouton density was subsequently quantified. No difference in total neurite number or length was observed using Sholl analysis (area under the curve 145±37.2 control vs 110±18.5 *IGFR-KO, p=0.51). Male *IGFR-KO mice did have fewer CA1 synaptic boutons than controls (**Figure 2B**, p=0.0002, n=9,12). Similar to the behavioral outcomes, no difference in synaptic bouton number was observed in female *IGFR-KOs (**Supplemental Figure 1D**, p=0.18, n=7,8). While the bouton number in the CA1 was significantly reduced in male mice, no significant differences in total protein levels of synaptophysin or PSD-95 were seen in full dorsal hippocampal lysates (**Supplemental Figure 1E-F**).

**Figure 2:**
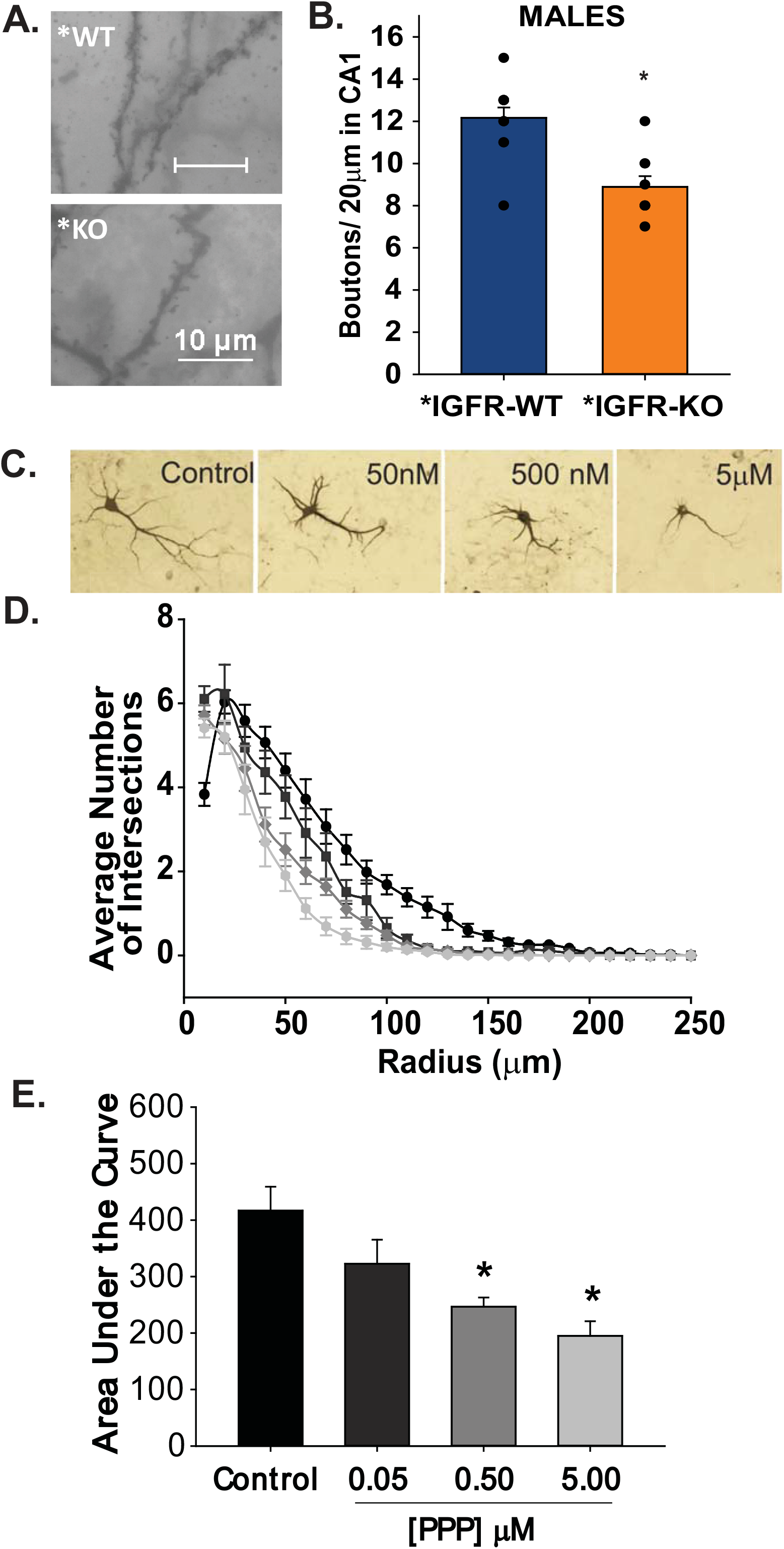
Alterations in hippocampal neuron structure with neuronal IGF-1R deficiency. (**A**) Representative images of synaptic boutons on dendrites in CA1 of *IGFR-WT and *IGFR-KO male mice. **(B)** The average number of boutons across 20μm of a dendrite within the CA1 (minimum 2 dendrites per mouse, mouse n=9,12). The asterisk denotes p<0.05, t-test. **(C)** Representative images of cultured hippocampal neurons treated with increasing concentrations of IGF-1R inhibitor, PPP. **(D)** Sholl analysis tracing quantification of the average number of crossings every 10 microns from the soma with increasing PPP, 24 hours following PPP administration. **(E)** The average area under the curve from **(D)**; (n=6,8 fields). All data are represented as mean +/- SEM and an asterisk denotes p< 0.05 vs the DMSO vehicle control (One-way ANOVA, post-hoc Dunnett’s test). *PPP* picropodophyllin toxin.

Changes in neuronal structure could also be noted in 8-10-day old primary hippocampal neuron cultures when IGFR signaling was reduced. Increasing concentrations of the small molecule IGFR inhibitor, picropodophyllin toxin (PPP), led to a concentration-dependent decrease in neurite complexity (**Figure 2C-E**). Note, neuron cultures were not stratified by sex as tissues were combined during isolation. Sholl analysis revealed a significant reduction in neurite length and complexity with 0.50μM-5μM PPP, compared to DMSO control (**Figure 2E-F**, p<0.05, n=8). Concentrations above 5μM were neurotoxic.

### Reduced MAPK and Increased ROCK with Neuronal IGFR Deficiency

Because male mice exhibited behavioral and cellular alterations when neuronal IGF-1R was reduced, the activation of downstream kinase signaling cascades within the hippocampal tissue was then assessed. As anticipated, phosphorylated MAPK was significantly reduced in *IGFR-KO hippocampi (**Figure 3A**, p= 0.009, n=8,12). Conversely, ROCK activity was significantly increased (**Figure 3B**, p=0.025, n=6,7). Downstream of ROCK, the ratio of phosphocofilin to total cofilin of *IGFR-KOs was trending but did not reach significance (**Figure 3C**, p=0.08, n= 7,8).

**Figure 3:**
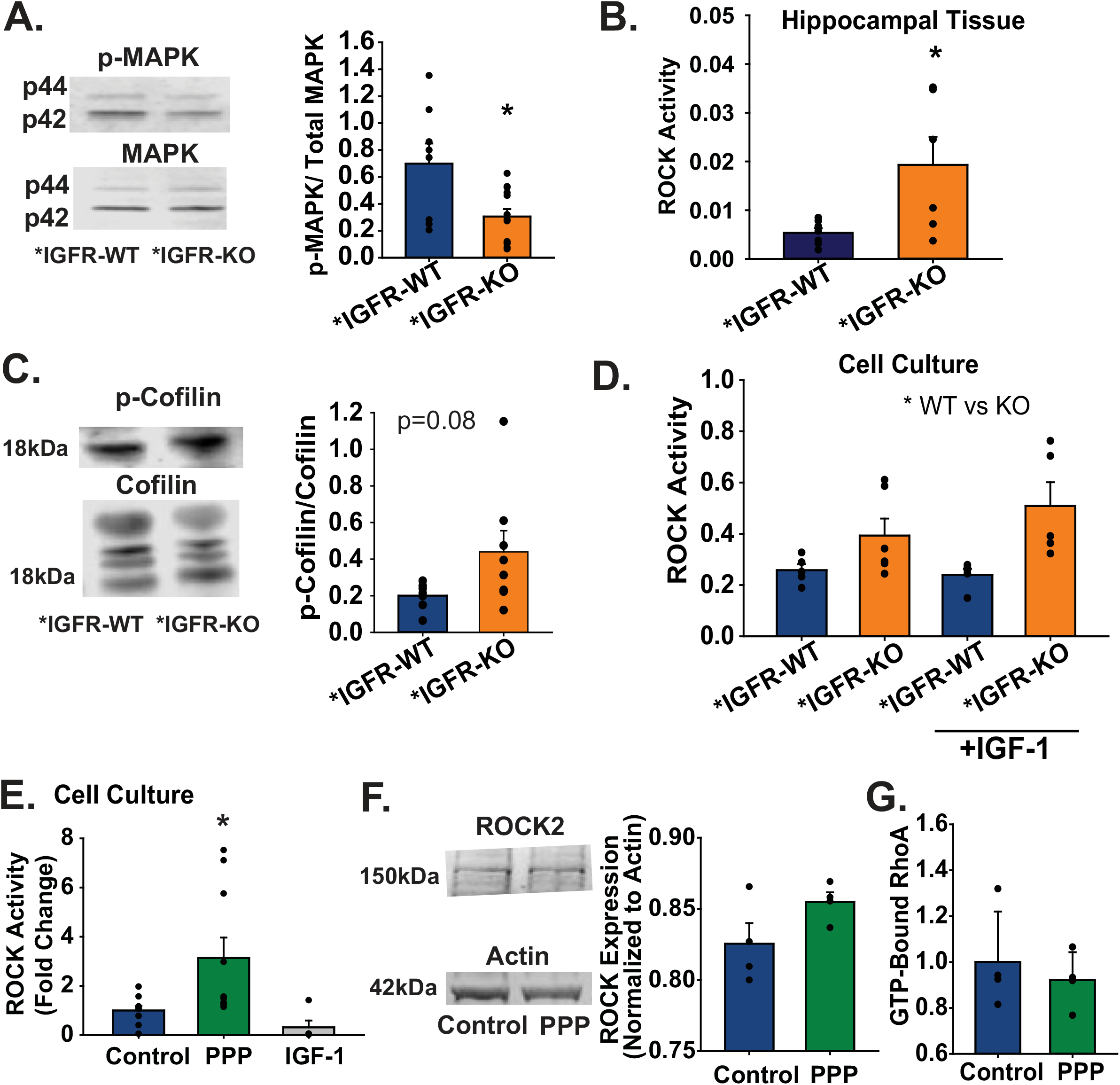
Alterations in MAPK and ROCK signaling with neuronal IGFR deficiency. **(A)** Left: Representative image of MAPK western blots, Right: Average ratio of phosphorylated MAPK to total MAPK in male *IGFR-WT and *IGFR-KO hippocampal lysates. The asterisk denotes p<0.05, t-test. **(B)** Average ROCK activity in male *IGFR-WT and *IGFR-KO hippocampal lysates. The asterisk denotes p<0.05, t-test. **(C)** Left: Representative image of western blots Right: Average ratio of phosphocofilin to total cofilin in male*IGFR-WT and *IGFR- KO hippocampal lysates. **(D)** ROCK activity in cultured *igfr* ^*f/f*^ neurons treated 48 hours after treatment with AAV9-Syn-Cre (or AAV9-Syn-GFP), with and without 100nM supplemented IGF-1 in cell culture. The asterisk indicates the significant main effect of WT vs KO, with no difference in IGF-1 treatment condition (two-way ANOVA posthoc Bonferonni, genetic treatment, and IGF-1 treatment as factors). **(E)** Average ROCK activity levels in cultured neurons treated with 0.5μM PPP, 100nM DMSO vehicle. The asterisk indicates a significant difference compared to DMSO control (One-way ANOVA, post-hoc Dunnett’s vs control). Average ROCK2 protein expression (Left: representative western blots, Right: quantification) **(F)** and GTP-bound RhoA levels **(G)** in cultured neurons treated with IGFR inhibitor 0.5μM PPP. All data is represented by mean +/- SEM.

Genetic reduction of IGF-1R in cultured neurons resulted in increased ROCK activity in the presence of exogenous IGF-1 in the media (**Figure 3D**, n=5-6). A two-way ANOVA comparison revealed significant main effects with genetic treatments (*IGFR-WT vs *IGFR-KO, p=0.004), and no effect from IGF-1 treatment (p=0.433). ROCK activity was also increased when cultured neurons were inhibited with 0.50μM PPP (**Figure 3E**, p=0.021, n=10). Supplemental IGF-1 did not significantly alter ROCK activity (**Figure 3E**, p=0.705). Mechanistically, neither PPP nor genetic reductions in IGF-1R lead to significant increases in ROCK2 protein expression in cultured neurons (**Figure 3F** p=0.16, **Supplemental Figure 2** p=0.117). mRNA levels of ROCK1, ROCK2, and upstream RhoA were unaffected by 0.50μM PPP (**Supplemental Figure 2A**, p>0.05). Neither were total RhoA protein expression, phosphorylated RhoA levels, or the levels of GTP-bound RhoA (**Supplemental Figure 2C** and **Figure 3G**, p>0.05), indicating the increase in ROCK activity was not due to robust changes in upstream RhoA activity.

### Restoration of Neurite Outgrowth with ROCK Inhibition

While it is difficult to pharmacologically increase MAPK activity, small molecule inhibitors of ROCK are readily available. Thus, neuron cultures were treated with small molecule ROCK inhibitors, Hydroxyfasudil (HF) and 1-Y27632, in an attempt to restore neurite outgrowth in the presence of the IGFR inhibitor PPP. Both 5μM HF and 1μM 1-Y27632 significantly increased neurite complexity on their own (**Figure 4A-C**, p<0.05, n=6) and prevented decreased complexity with PPP **(Figure 4A-C**, p<0.05). No significant difference was observed in the rescued growth between the either pharmacological ROCK inhibitors.

**Figure 4:**
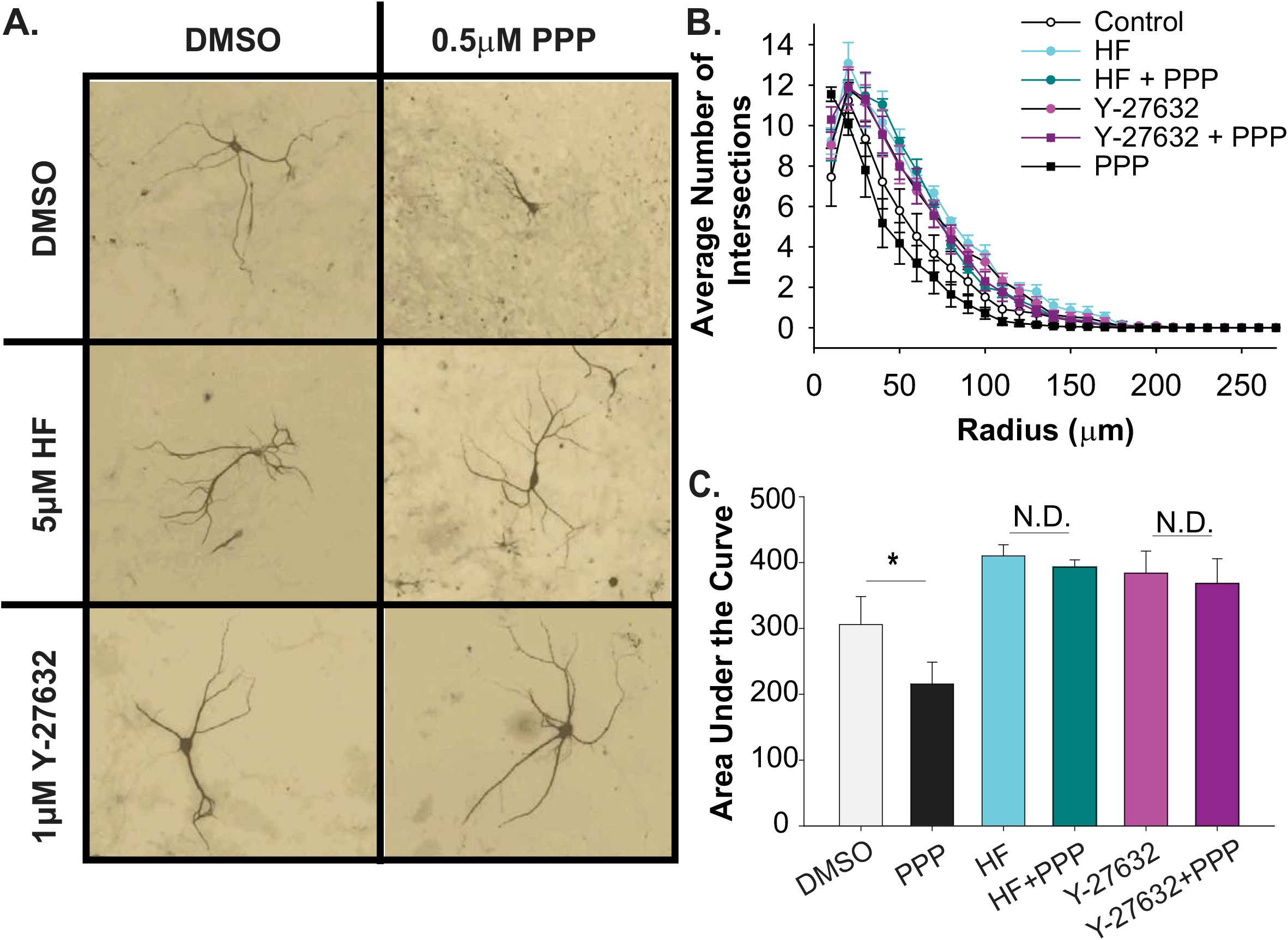
The pharmacological inhibition and restoration of neurite outgrowth with IGF-1R and ROCK inhibitors. **(A)** Representative images of neuronal growth with PPP (IGF-1R inhibitor) and ROCK inhibitors (HF and Y-27632). **(B)** Sholl analysis of the average number of neuronal crossings every 10μm from the soma in neurons treated with DMSO vehicle, 0.5μM PPP, 5μM HF, and 1μM Y-27632 treatments. **(C)** The average area under the curves in **B** (6 neurons per coverslip, n=3,4 coverslips). Asterisk denotes p<0.05 for knock-out effect, (Two-way ANOVA, post-hoc Bonferonni multiple comparisons). Abbreviations: *N*.*D*. no difference, *PPP* picropodophyllin toxin, *HF* Hydroxyfasudil.

### Spatial Learning and Neuronal Structure Restoration with ROCK Inhibition

Based on the structural restoration *in vitro*, we administered 10 mg/kg HF to male *IGFR-KO mice via osmotic pump for 14 days and assessed whether spatial learning and memory deficits could also be rescued (**Figure 5A-B** and **Supplemental Figure 2**). Spatial learning deficits of male *IGFR-KOs in the acquisition phase of the radial arm water maze were not restored after receiving 14-days of HF infusion (**Figure 5A-B**, p>0.05, n=14). In the reversal phase, HF treated *IGFR-KO mice were no different than *IGFR-KO-DMSO controls nor were they statistically different than *IGFR-WT-HF, suggesting a possible partial rescue (**Figure 5B**). Two-way ANOVA comparisons revealed significant main effects from genetic treatment groups (p=0.034), with no pharmacological drug treatment effect (p=0.503). The total number of errors to find the escape platform in the acquisition and reversal phases showed no significant effects of HF treatment in the *IGFR-WT and/or *IGFR-KO mice (**Supplemental Figure 3**).

**Figure 5:**
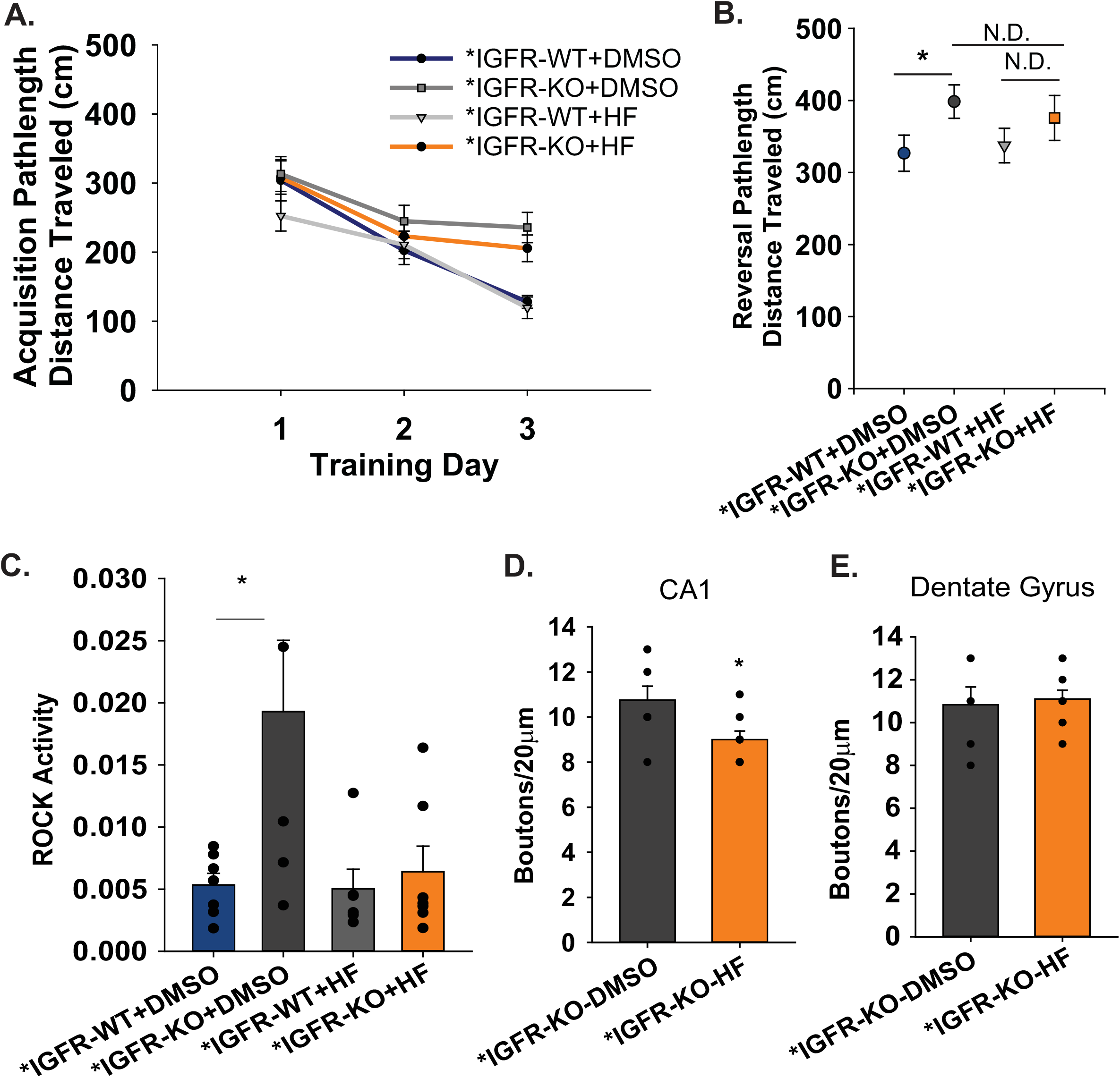
Partial restoration of spatial learning and memory and synaptic bouton number with HF infusion. Mice were implanted with 14-day releasing mini-osmotic pumps loaded with 10mg/kg HF or DMSO vehicle. **(A)** Average total pathlength traveled per trial on each day of the acquisition phase of the radial arm water maze (n=14). **(B)** Average pathlength traveled of four treatment groups during the reversal phase of the radial arm water maze (n=14). Asterisk denotes p<0.05 for WT vs KO, while N.D. denotes no statistical difference (p>0.05, two-way ANOVA, post-hoc Bonferonni multiple comparisons) **(C)** Average ROCK activity within the hippocampal lysates of *IGFR-WT and *IGFR-KO mice treated with DMSO vehicle and HF (n=6,7). Average number of synaptic boutons across 20μm of dendrites in the CA1 (**D**) and DG (**E**) (n=7,10). Asterisk denotes p<0.05 to respective control (t-test). All data are expressed as mean +/- SEM.

Immediately following behavioral analysis, hippocampal tissue was isolated to assess ROCK activity. A significant increase in ROCK activity was noted in the hippocampi of *IGFR-KO-DMSO mice compared to *IGFR-WT-DMSO controls (**Figure 5C**, p=0.02, n=6,7). No statistical difference was noted in *IGFR-KO-HF mice, suggesting bioavailability and accumulation of the ROCK inhibitor in the hippocampus (**Figure 5C**, p>0.05, n=7). However, Golgi-Cox staining revealed that HF treatment did not increase the number of boutons in the CA1 or dentate gyrus, in fact, the bouton number was significantly decreased in the CA1 of mice receiving the ROCK inhibitor (**Figure 5D-E**, p=0.03, n=8).

## DISCUSSION

The data presented here demonstrate that neuronal IGF-1 signaling is necessary to maintain spatial learning and memory in adult male mice. Previous studies have highlighted how developmental reductions in IGF-1 lead to impairments in neuroglial function, and ultimately impairments in cognitive function. Genetic knockouts of IGF-1 and IGF-1R exhibit reduced brain size decreased myelination, and several behavioral deficits (Beck et al., 1995). The protective effects of IGF-1 on cognitive function are not limited to early-life phenotypes, as the natural decline in circulating IGF-1 in advanced age is associated with increased cognitive deficits in both humans and animal models (Ashpole et al., 2015b; Deijen et al., 2011; Deijen et al., 1998; van Dam et al., 2000). Moreover, the age-related loss of circulating IGF-1 is thought to predispose the aged brain to Alzheimer’s Disease and related dementias (Deak and Sonntag, 2012; Doi et al., 2015; Licht et al., 2014; Sonntag et al., 2013; Vidal et al., 2016). Unfortunately, animal models of reduced circulating IGF-1 also exhibit alterations in cardiovascular function, blood-brain barrier integrity, muscle strength, and bone structure due to the extensive anabolic effects of IGF-1R throughout the body (Ashpole et al., 2016; Ashpole et al., 2015a; Ashpole et al., 2017; Farias Quipildor et al., 2019; Kinney et al., 2001; Tarantini et al., 2016a; Tarantini et al., 2016b). This underscores how systemic changes in IGF-1 induces numerous variables that may hamper our interpretation of underlying mechanisms. Thus, we aimed to utilize a mouse model where IGF-1 signaling could be reduced in tissue and cell-specific manner. In addition, we wanted to control the timing of the reduction in IGF-1 signaling to avoid developmental confounds of possible compensatory increases in local IGF-1 production, insulin receptor activity, and insulin-like binding protein prevalence. Therefore, we aimed to delineate the role of neuronal IGF-1 signaling on spatial and working learning and memory in adulthood, after the pubertal surge in IGF-1 has peaked. In the present study, neuronal reduction of IGF-1 in the hippocampus of male mice led to impaired spatial learning, reduced synaptic bouton number, and increased ROCK signaling. While ROCK inhibition rescued neurite outgrowth in vitro, it did not fully restore learning and memory in vivo.

Impairments in spatial learning and memory in the neuronal IGFR-KO mice were accompanied by reductions in hippocampal synapse number, highlighting the possibility that synapse retraction was a potential mechanism underlying cognitive dysfunction. Synaptic pruning and restructuring are markedly increased during puberty, particularly within the hippocampal regions of interest (Anderson et al., 2002). Our goal was to reduce neuronal IGF-1R after this critical developmental window; however, we cannot differentiate whether the reduction in synaptic boutons observed in our knock-out mice is due to increased synaptic pruning or synapse retraction. The possibility of synaptic pruning within our mutant mice is supported by evidence that synaptic pruning is activity-dependent, and IGF-1 significantly increases synaptic excitability by modulating NMDA receptor availability, calcium channel function, and inhibitory neurotransmission (Blair et al., 1999; Cao et al., 2011; Gazit et al., 2016; Le Greves et al., 2005; Mardinly et al., 2016). While the reductions in IGF-1R were restricted to neurons, in future studies it would be important to assess microglial activation and phagocytosis within the hippocampus of our neuronal IGFR-KO mice to assess whether this mutation increased synaptic pruning. Further analyses of total neuron number and/or newborn neuron incorporation and survival may yield additional information on cellular changes with neuronal IGF-R knockout, especially when one considers previous evidence that IGF-1 promotes survival and differentiation of neurons (Brooker et al., 2000; D’Mello et al., 1997; Nieto-Estevez et al., 2016). Moreover, a detailed analysis of synapse structure may highlight additional structural differences, as IGF-1 supplementation in aged rats has been shown to increase post-synaptic density and the number of multiple spine boutons(Shi et al., 2005).

Considering the localized restriction of neuronal IGFR deficiency in our study, we focused on hippocampal-dependent behaviors as read-outs of cognitive function. The radial arm water maze showed robust between-group and over-time differences. However, differences were not observed in the novelty recognition assay. The lack of treatment effect within this assay may be influenced by the maintenance of IGF-1 signaling in the other brain regions implicated in novelty seeking behaviors. On the other hand, neither treatment group showed robust novelty discrimination scores above 0.7. Inclusion of wild-type controls without stereotactic injection of control vectors may be of use to rule out confounds of superior tissue displacement and damage. Additionally, future inducible models of IGFR deletion throughout the entire adult brain may be of interest, as a recent study highlighted that overexpression of IGF-1 within the aged brain had behavior-specific outcomes (Farias Quipildor et al., 2019).

Reductions in MAPK, ERK, and Akt signaling were anticipated with IGF-1 deficiency, as these are the canonical pathways downstream of IGF-1R activation. In addition to reduced MAPK within the hippocampus of our neuronal IGFR-KO mice, we also observed a significant upregulation of RhoA/ROCK activity. As this pathway is well-known to decrease neuronal excitability and induce spine retraction, the observed increase in our neuronal IGFR-KO mice supported our working hypothesis that reduced bouton number may be due to synapse retraction. We utilized a series of in vitro studies to further examine the relationship between IGF-1 and ROCK signaling. Consistent with previous studies, significant increases in neurite extension were observed when RhoA/ROCK signaling was reduced (Chen and Firestein, 2007; Greathouse et al., 2018). Furthermore, ROCK inhibition circumvented the stunted growth associated with IGF-1 inhibition, suggesting a possible route of rescue in our mutant mice. This was particularly interesting considering previous studies have shown small molecule inhibitors of ROCK, including HF, can rescue spatial learning and memory in aged mice and rodent models of neurodegeneration (Guo et al., 2020; Huentelman et al., 2009; Yu et al., 2017). Unfortunately, we did not observe a rescue of synapse number or learning and memory despite our doses of HF falling within the range of these previous studies. In fact, HF mice perplexingly displayed fewer synaptic boutons than controls, without showing signs of spatial learning impairment. It is unclear why bouton number was lower in the HF-treated mice, but it was apparent that ROCK was inhibited in this tissue, which indicates the inhibitor was available when tissues were isolated on the last day of drug administration. When higher-order cognitive flexibility was assessed, the HF knock-outs were not different than wild-type controls or knock-out controls, possibly suggesting a partial rescue. However, this phenotype was not robust nor was it prevalent when other endpoints were examined. It is possible that alternative outcomes would have been observed had ROCK been genetically targeted at the same time neuronal IGF-1R was reduced. ROCK inhibitors were not administered until two weeks before behavioral assessment (6 weeks following IGF-1R reduction). Although the 14-day treatment is sufficient to reduce ROCK activity levels to those of controls, it may not be sufficient to allow the reversal of cognitive deficits. Indeed, many of the previous studies with ROCK inhibitors dose animals for two or more months (Busti et al., 2020; Takata et al., 2013; Tonges et al., 2012; Yu et al., 2017). A prophylactic approach may be advantageous to solidify the contributions of ROCK to IGF-1 related deficits.

Sex-specific differences in the response to IGF-1 are increasingly reported (Ashpole et al., 2017; Austad and Fischer, 2016; Bokov et al., 2011; Farias Quipildor et al., 2019; Mao et al., 2018), so it was not surprising to observe impairments limited to male mice. Of note, our experimental design was set for independent assessments of male and female cohorts, rather than cross-comparisons. While sex-specific differences are commonly observed in aged, IGF-1 deficient animals, the mechanism underlying these disparities is not understood. A role for sex hormones has been proposed (Austad, 2019; Austad and Fischer, 2016); however, concrete evidence for this is still lacking. Additional studies focused on the mechanism underlying the sex-dependence of the effect of IGF-1 in the brain are warranted, and would reveal critical information needed across the fields of developmental biology and geroscience.

Overall, we observe significant impairments in spatial learning and memory when IGF-1 signaling is reduced in neurons of adult male mice. These results provide potential mechanistic insights underlying learning and memory impairments in advanced age when IGF-1 levels are significantly reduced. Considering the therapeutic limitations of exogenous IGF-1 administration in the elderly, the new focus should now turn downstream of neuronal IGF-1R to identify potential therapeutics that could rescue or prevent the loss of cognitive function.

## Acknowledgments

The authors would like to thank the animal care staff at both the University of Oklahoma Health Sciences Center and the University of Mississippi.

## Supplemental Information- Hayes et al

**Supplemental Figure 1:**
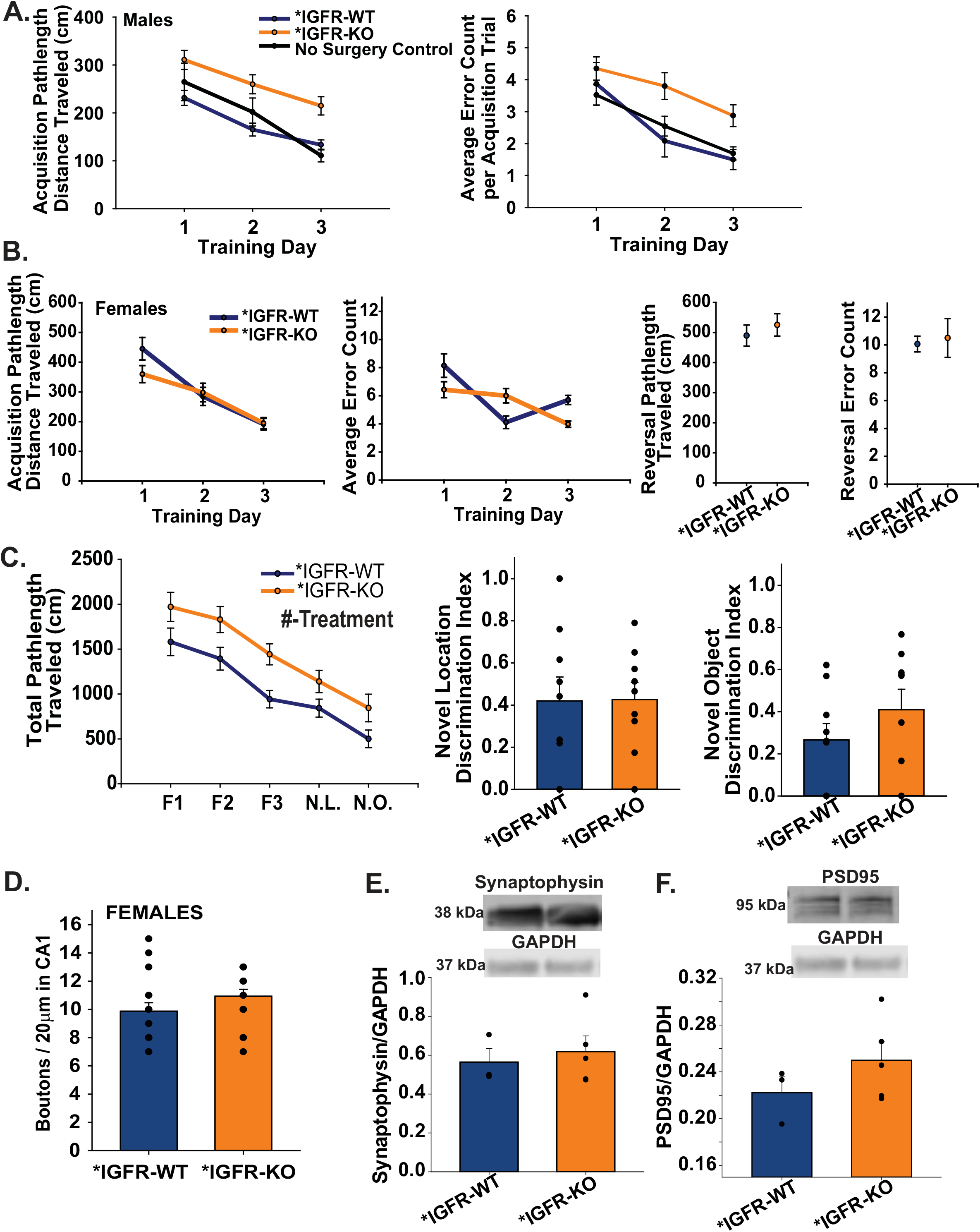
Sex-specific alterations in learning and memory and neuron structure with hippocampal neuron IGFR deficiency. (**A**) Average pathlength traveled and errors made during radial arm maze acquisition in non-surgical control, surgical control (*IGFR-WT), and surgical knockout (*IGFR-KO) male mice. (**B)** Average pathlength traveled and errors made by female mice during the acquisition and reversal phase of radial arm water maze (n= 9,10). **(C)** Working memory, novel location discrimination index, and novel object discrimination index in female *IGFR-WT and *IGFR-KO (n=9,10). **(D)** The average of synaptic boutons across 20 microns of dedrites in the CA of female *IGFR-WT and *IGFR-KO (n=8,8). **(E)** Top: Representative images of syanptophysin and GAPDH western blots; bottom the average ratio of sypnatophysin to GAPDH protein expression in male *IGFR-WT and *IGFR-KO (n=3,5). **(F)** Top: representative image of PSD95 and GAPDH western blots; bottom: the average ratio of PSD95 to GAPDH protein expression in female *IGFR-WT and *IGFR-KO hippocampal tissue (n=3,5). #Treatment denotes a difference between groups (p<0.05; Two-way ANOVA repeated measures). All data represent mean +/- SEM. Abbreviations: *F1,F2,F3* indicates Familiarization trial number. *N*.*L* novel location, *N*.*O*. novel object.

**Supplemental Figure 2:**
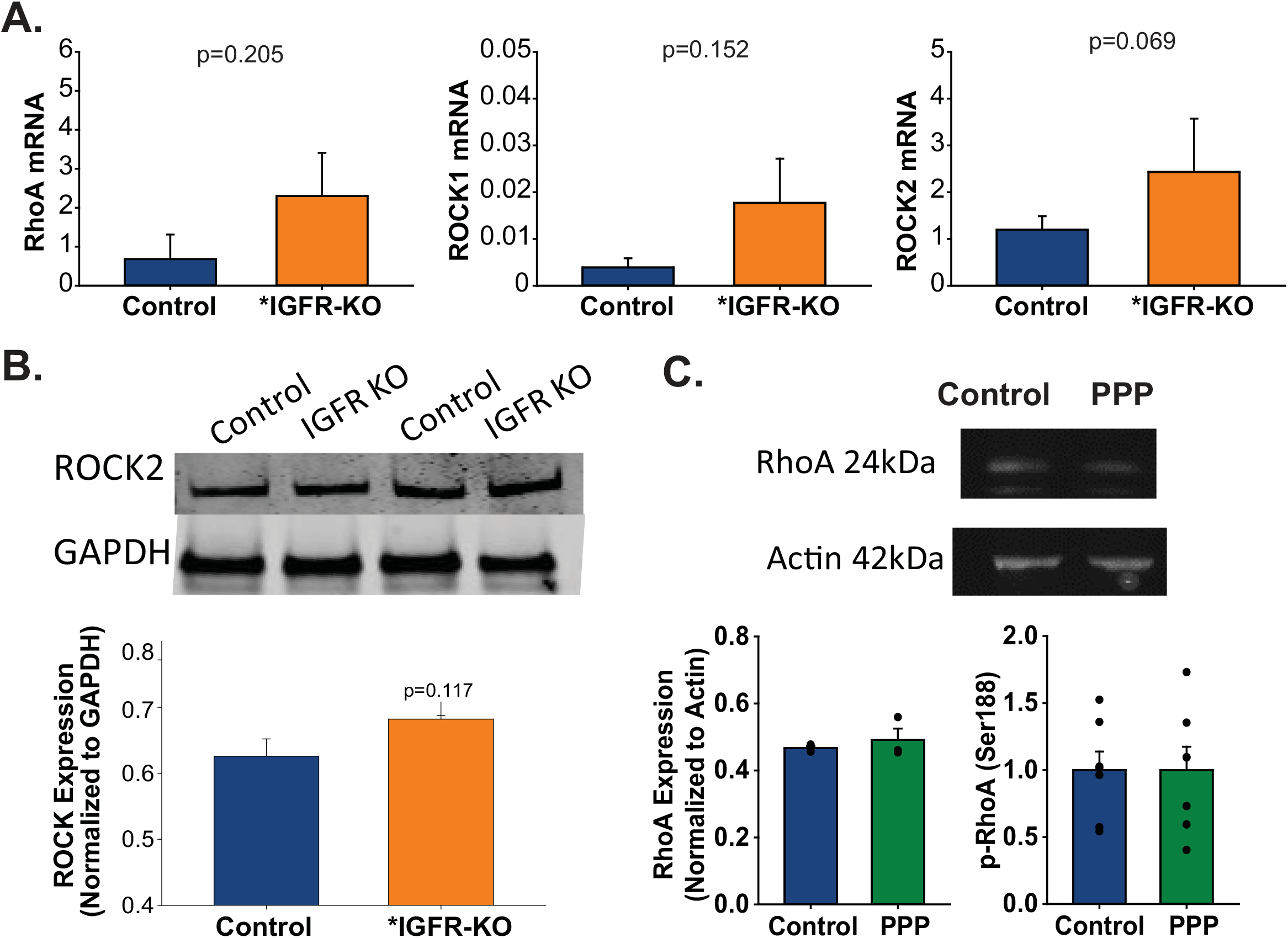
**(A)** Average mRNA expression of RhoA, ROCK1, and ROCK2 in neurons deficient in IGF-1R (AAV-mediated Cre recombinase in *igfr*^*f/f*^ neurons). **(B)** Average ROCK2 protein expression in hippocampal neurons deficient in IGF-1R. **(C)** Top: Representative image of RhoA western blot, Bottom: Average RhoA and phosphorylated RhoA levels (Ser188) in cultured neurons treated with 0.5μM PPP or DMSO control.

**Supplemental Figure 3:**
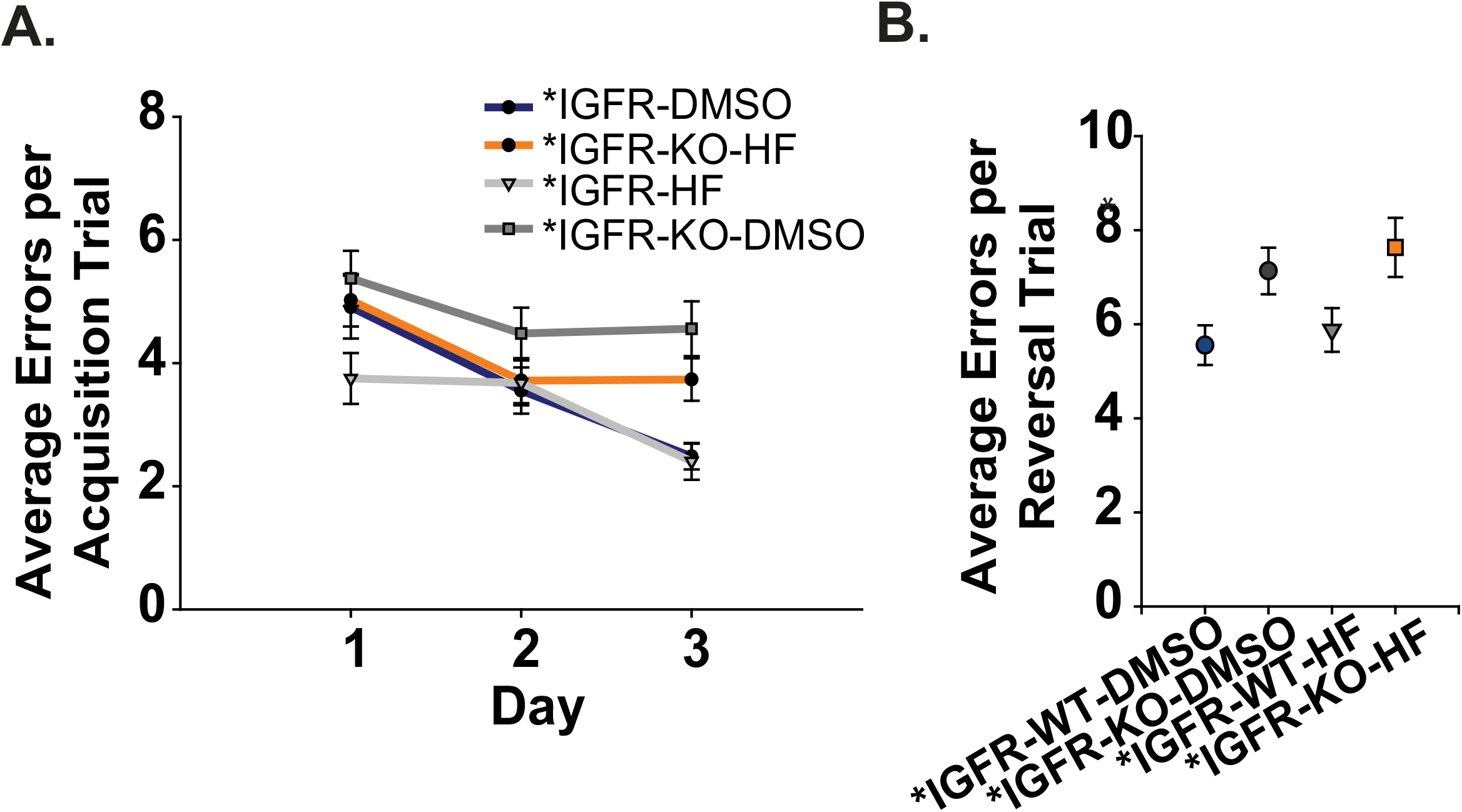
Spatial learning and memory in RAWM of males that received pharmacological treatments. **(A)** The average total number of errors between *IGFR-WT and *IGFR-KO that received DMSO and HF in the acquisition phase of RAWM (n=14). **(B)** The average total number of errors in the reversal phase of RAWM between four treatment groups (n=14). **(C)** The average number of boutons on dendrites in the dentate gyrus of the hippocampus in *IGFR-KOs that received treatments, DMSO and HF (n=7,10). All data are represented as mean +/- SEM. Asterisk denotes p<0.05 vs. respective control (t-test).

**Supplemental Table 1:**
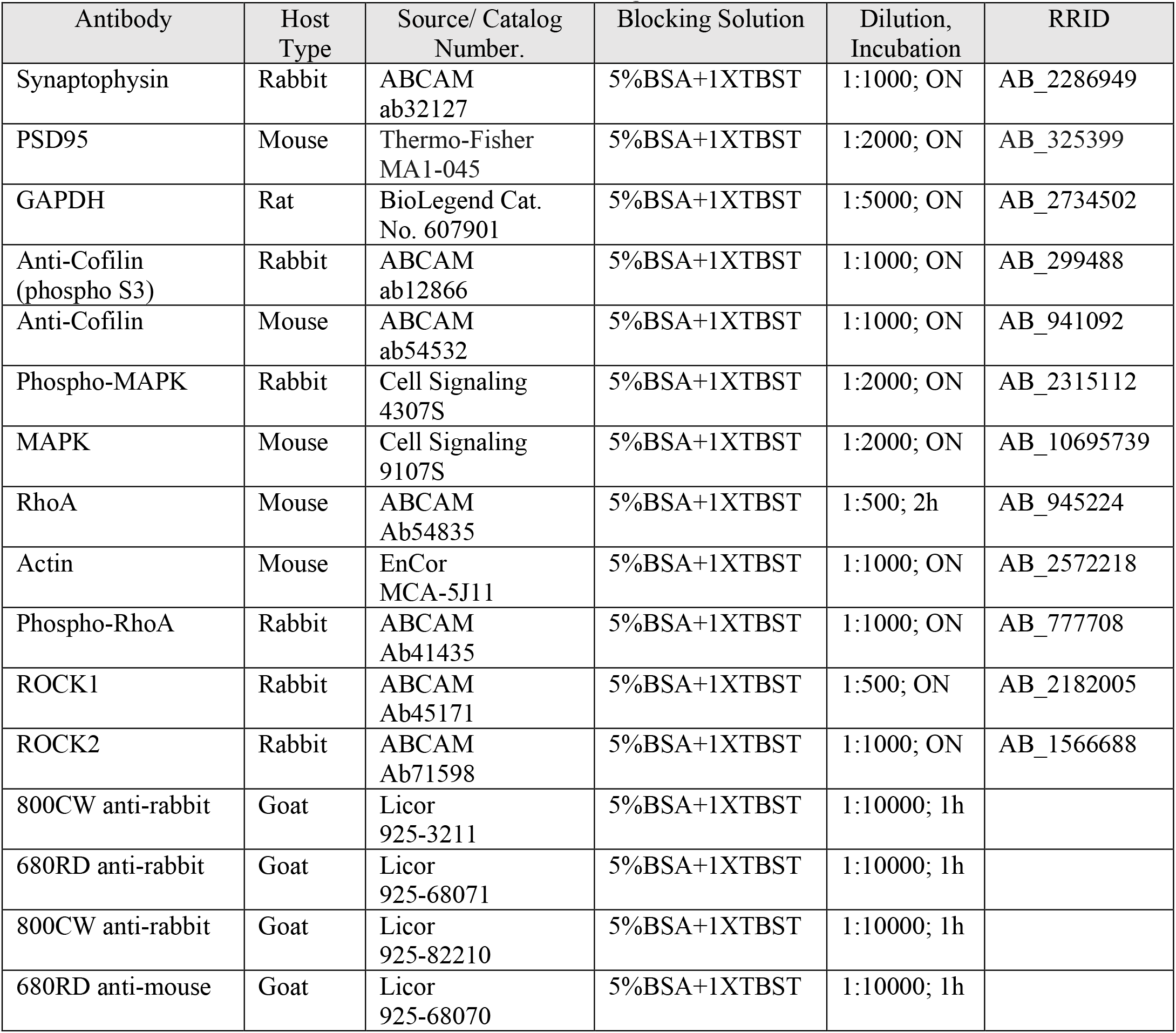
Antibodies used with RRIDs. Abbreviations: *BSA* bovine serum albumin, *TBST* Tris Buffered Saline with Tween, *ON* overnight, *h* hour

**Supplemental Table 2:**
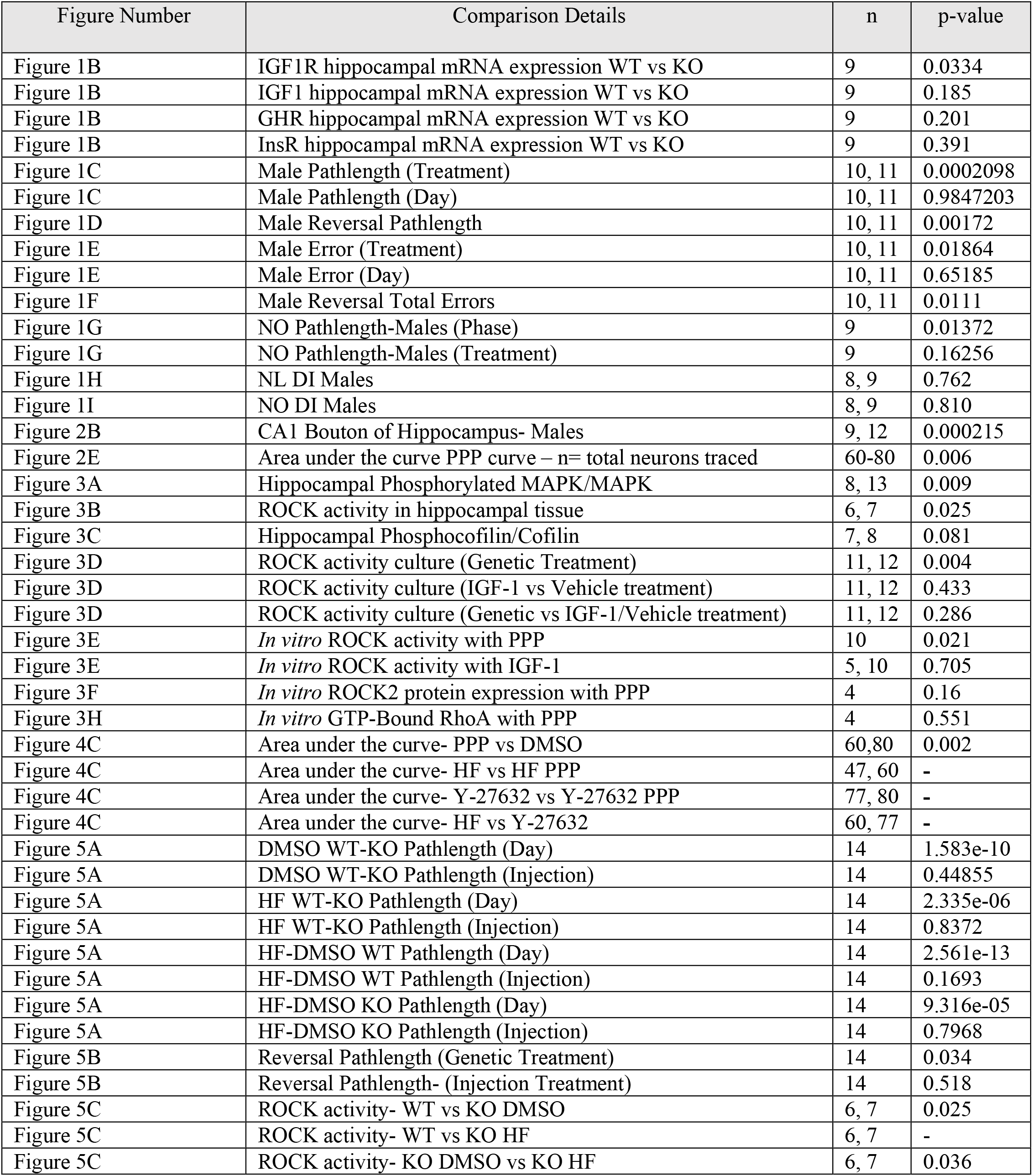
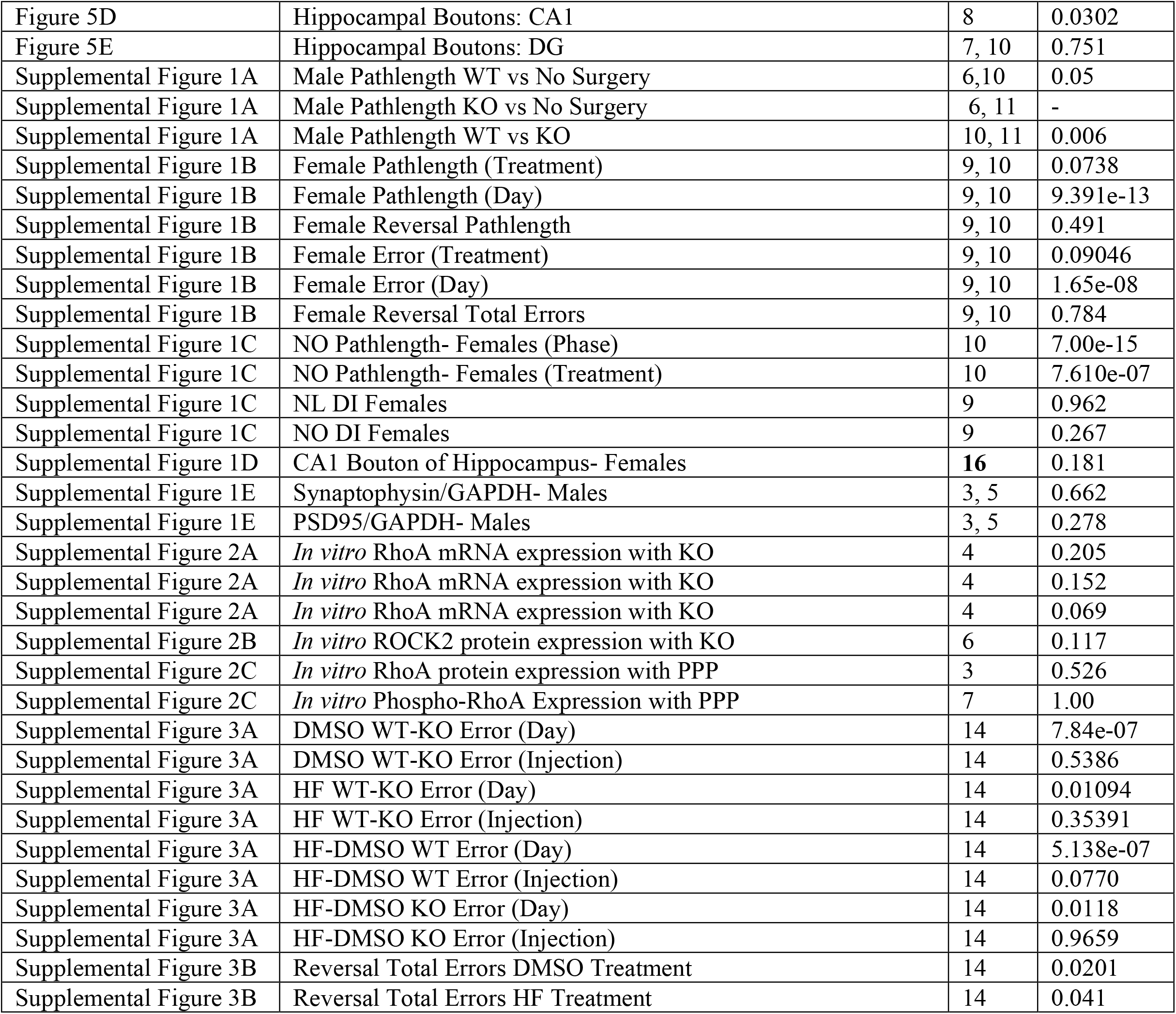
A summarized table of statistical comparisons denoting figure, details of comparisons, sample size, and P values. **Abbreviations:** *HF* Hydroxyfasudil, *PPP* picropodophyllin toxin, *DMSO* dimethylsulfoxide, *WT* *IGFR-WT wildtype control that received AAV9-Syn-GFP, *KO* knockout of neuronal IGF-1R (*IGFR-KO) that received AV9-Syn-Cre, *NL* Novel Location, *NO* Novel Object, *DI* discrimination index, *GHR* growth hormone receptor, *InsR* Insulin Receptor, *IGF1* Insulin-Like Growth Factor-1, *IGFR* Insulin-like growth factor receptor, *ROCK* Rho Kinase

| Figure Number | Comparison Details | n | p-value |
| --- | --- | --- | --- |
| Figure 1B | IGF1R hippocampal mRNA expression WT vs KO | 9 | 0.0334 |
| Figure 1B | IGF1 hippocampal mRNA expression WT vs KO | 9 | 0.185 |
| Figure 1B | GHR hippocampal mRNA expression WT vs KO | 9 | 0.201 |
| Figure 1B | InsR hippocampal mRNA expression WT vs KO | 9 | 0.391 |
| Figure 1C | Male Pathlength (Treatment) | 10, 11 | 0.0002098 |
| Figure 1C | Male Pathlength (Day) | 10, 11 | 0.9847203 |
| Figure 1D | Male Reversal Pathlength | 10, 11 | 0.00172 |
| Figure 1E | Male Error (Treatment) | 10, 11 | 0.01864 |
| Figure 1E | Male Error (Day) | 10, 11 | 0.65185 |
| Figure 1F | Male Reversal Total Errors | 10, 11 | 0.0111 |
| Figure 1G | NO Pathlength-Males (Phase) | 9 | 0.01372 |
| Figure 1G | NO Pathlength-Males (Treatment) | 9 | 0.16256 |
| Figure 1H | NL DI Males | 8, 9 | 0.762 |
| Figure 1I | NO DI Males | 8, 9 | 0.810 |
| Figure 2B | CA1 Bouton of Hippocampus- Males | 9, 12 | 0.000215 |
| Figure 2E | Area under the curve PPP curve – n= total neurons traced | 60-80 | 0.006 |
| Figure 3A | Hippocampal Phosphorylated MAPK/MAPK | 8, 13 | 0.009 |
| Figure 3B | ROCK activity in hippocampal tissue | 6, 7 | 0.025 |
| Figure 3C | Hippocampal Phosphocofilin/Cofilin | 7, 8 | 0.081 |
| Figure 3D | ROCK activity culture (Genetic Treatment) | 11, 12 | 0.004 |
| Figure 3D | ROCK activity culture (IGF-1 vs Vehicle treatment) | 11, 12 | 0.433 |
| Figure 3D | ROCK activity culture (Genetic vs IGF-1/Vehicle treatment) | 11, 12 | 0.286 |
| Figure 3E | *In vitro* ROCK activity with PPP | 10 | 0.021 |
| Figure 3E | *In vitro* ROCK activity with IGF-1 | 5, 10 | 0.705 |
| Figure 3F | *In vitro* ROCK2 protein expression with PPP | 4 | 0.16 |
| Figure 3H | *In vitro* GTP-Bound RhoA with PPP | 4 | 0.551 |
| Figure 4C | Area under the curve- PPP vs DMSO | 60,80 | 0.002 |
| Figure 4C | Area under the curve- HF vs HF PPP | 47, 60 | - |
| Figure 4C | Area under the curve- Y-27632 vs Y-27632 PPP | 77, 80 | - |
| Figure 4C | Area under the curve- HF vs Y-27632 | 60, 77 | - |
| Figure 5A | DMSO WT-KO Pathlength (Day) | 14 | 1.583e-10 |
| Figure 5A | DMSO WT-KO Pathlength (Injection) | 14 | 0.44855 |
| Figure 5A | HF WT-KO Pathlength (Day) | 14 | 2.335e-06 |
| Figure 5A | HF WT-KO Pathlength (Injection) | 14 | 0.8372 |
| Figure 5A | HF-DMSO WT Pathlength (Day) | 14 | 2.561e-13 |
| Figure 5A | HF-DMSO WT Pathlength (Injection) | 14 | 0.1693 |
| Figure 5A | HF-DMSO KO Pathlength (Day) | 14 | 9.316e-05 |
| Figure 5A | HF-DMSO KO Pathlength (Injection) | 14 | 0.7968 |
| Figure 5B | Reversal Pathlength (Genetic Treatment) | 14 | 0.034 |
| Figure 5B | Reversal Pathlength- (Injection Treatment) | 14 | 0.518 |
| Figure 5C | ROCK activity- WT vs KO DMSO | 6, 7 | 0.025 |
| Figure 5C | ROCK activity- WT vs KO HF | 6, 7 | - |
| Figure 5C | ROCK activity- KO DMSO vs KO HF | 6, 7 | 0.036 |
| Figure 5D | Hippocampal Boutons: CA1 | 8 | 0.0302 |
| Figure 5E | Hippocampal Boutons: DG | 7, 10 | 0.751 |
| Supplemental Figure 1A | Male Pathlength WT vs No Surgery | 6,10 | 0.05 |
| Supplemental Figure 1A | Male Pathlength KO vs No Surgery | 6, 11 | - |
| Supplemental Figure 1A | Male Pathlength WT vs KO | 10, 11 | 0.006 |
| Supplemental Figure 1B | Female Pathlength (Treatment) | 9, 10 | 0.0738 |
| Supplemental Figure 1B | Female Pathlength (Day) | 9, 10 | 9.391e-13 |
| Supplemental Figure 1B | Female Reversal Pathlength | 9, 10 | 0.491 |
| Supplemental Figure 1B | Female Error (Treatment) | 9, 10 | 0.09046 |
| Supplemental Figure 1B | Female Error (Day) | 9, 10 | 1.65e-08 |
| Supplemental Figure 1B | Female Reversal Total Errors | 9, 10 | 0.784 |
| Supplemental Figure 1C | NO Pathlength- Females (Phase) | 10 | 7.00e-15 |
| Supplemental Figure 1C | NO Pathlength- Females (Treatment) | 10 | 7.610e-07 |
| Supplemental Figure 1C | NL DI Females | 9 | 0.962 |
| Supplemental Figure 1C | NO DI Females | 9 | 0.267 |
| Supplemental Figure 1D | CA1 Bouton of Hippocampus- Females | **16** | 0.181 |
| Supplemental Figure 1E | Synaptophysin/GAPDH- Males | 3, 5 | 0.662 |
| Supplemental Figure 1E | PSD95/GAPDH- Males | 3, 5 | 0.278 |
| Supplemental Figure 2A | *In vitro* RhoA mRNA expression with KO | 4 | 0.205 |
| Supplemental Figure 2A | *In vitro* RhoA mRNA expression with KO | 4 | 0.152 |
| Supplemental Figure 2A | *In vitro* RhoA mRNA expression with KO | 4 | 0.069 |
| Supplemental Figure 2B | *In vitro* ROCK2 protein expression with KO | 6 | 0.117 |
| Supplemental Figure 2C | *In vitro* RhoA protein expression with PPP | 3 | 0.526 |
| Supplemental Figure 2C | *In vitro* Phospho-RhoA Expression with PPP | 7 | 1.00 |
| Supplemental Figure 3A | DMSO WT-KO Error (Day) | 14 | 7.84e-07 |
| Supplemental Figure 3A | DMSO WT-KO Error (Injection) | 14 | 0.5386 |
| Supplemental Figure 3A | HF WT-KO Error (Day) | 14 | 0.01094 |
| Supplemental Figure 3A | HF WT-KO Error (Injection) | 14 | 0.35391 |
| Supplemental Figure 3A | HF-DMSO WT Error (Day) | 14 | 5.138e-07 |
| Supplemental Figure 3A | HF-DMSO WT Error (Injection) | 14 | 0.0770 |
| Supplemental Figure 3A | HF-DMSO KO Error (Day) | 14 | 0.0118 |
| Supplemental Figure 3A | HF-DMSO KO Error (Injection) | 14 | 0.9659 |
| Supplemental Figure 3B | Reversal Total Errors DMSO Treatment | 14 | 0.0201 |
| Supplemental Figure 3B | Reversal Total Errors HF Treatment | 14 | 0.041 |

